# Ontology-based modeling, integration, and analysis of heterogeneous clinical, pathological, and molecular kidney data for precision medicine

**DOI:** 10.1101/2024.04.01.587658

**Authors:** Yongqun Oliver He, Laura Barisoni, Avi Z Rosenberg, Peter N. Robinson, Alexander D. Diehl, Yichao Chen, Jimmy P. Phuong, Jens Hansen, Bruce W. Herr, Katy Börner, Nikki Bonevich, Ghida Arnous, Saketh Boddapati, Jie Zheng, Fadhl Alakwaa, Pinaki Sardar, William D. Duncan, Chen Liang, M. Todd Valerius, Sanjay Jain, Ravi Iyengar, Jonathan Himmelfarb, Matthias Kretzler, the Kidney Precision Medicine Project

**Affiliations:** University of Michigan, Ann Arbor, MI, USA; Duke University, Durham, NC, USA; Johns Hopkins University School of Medicine, MD, USA; The Jackson Laboratory for Genomic Medicine, Farmington, CT, USA; University at Buffalo, Buffalo, NY, USA; Pennsylvania State University, State College, PA, USA; University of Washington, Seattle, WA, USA; Icahn School of Medicine at Mount Sinai, New York, NY, USA; Indiana University, Bloomington, IN, USA; University of Florida, Gainesville, FL, USA; University of South Carolina, Columbia, SC, USA; Brigham and Women’s Hospital, Boston, Massachusetts, USA; Washington University, St. Louis, MO, USA

## Abstract

Many resources are now generating, processing, storing, or providing kidney-related molecular, pathological, and clinical data. Reference ontologies offer an opportunity to support knowledge and data organization and integration. The Kidney Precision Medicine Project (KPMP) team contributed to the representation and addition of 329 kidney phenotype terms to the Human Phenotype Ontology (HPO) and identified many subcategories of acute kidney injury (AKI) or chronic kidney disease (CKD). The Kidney Tissue Atlas Ontology (KTAO) imports and integrates kidney-related terms from existing ontologies (e.g., HPO, CL, and Uberon) and represents 259 kidney-related biomarkers. We also developed a precision medicine metadata ontology (PMMO) to integrate 50 variables from KPMP and CellxGene resources and applied PMMO for integrative analysis. The gene expression profiles of kidney gene biomarkers were specifically analyzed in healthy controls or AKI/CKD disease states. This work demonstrates how ontology-based approaches support multi-domain data and knowledge organization and integration to advance precision medicine.

## Introduction

The Kidney Precision Medicine Project (KPMP) (https://www.kpmp.org/) is an NIH/NIDDK-funded consortium that aims to characterize with precision the complexity of chronic kidney disease (CKD) and acute kidney injury (AKI) at the patient level to increase our ability to improve personalized treatment (1). While AKI is a sudden and often temporary loss of kidney function, CKD causes reduced kidney function over a long period of time and can lead to end-stage kidney disease. However, a transition from AKI to CKD can also be observed. AKI and CKD are associated with a complex pathogenesis involving genetic, pathological, molecular, social, and environmental factors. Despite significant efforts, the mechanisms underlying the development and progression of kidney diseases are not yet fully understood, partly because of challenges to integrating data from multiple knowledge domains. Hence, integrating different types of data related to kidney diseases deserves to be the subject of further in-depth investigations.

Significant effort has been made recently to generate kidney-related data and make them publicly available to the community of investigators. The Human BioMolecular Atlas Program (HuBMAP) aims to develop an open and global platform to map healthy cells in the human body (2). The Human Cell Atlas project (HCA) is a global consortium focused on mapping every cell type in the human body and developing a 3-dimensional atlas of human cells to transform our understanding of biology and disease (3). The CellxGene resource, which is closely associated with the HCA, is a suite of computational tools that help scientists deposit, download, query, and visually explore curated and standardized single-cell biology datasets (4).

Data integration remains a big challenge due to data heterogeneity within and between domains. Interoperable ontologies support data standardization, sharing, and integration. Many reference biomedical ontologies such as Cell Ontology (CL) (5), Uber-anatomy ontology (Uberon) (6), and Human Phenotype Ontology (HPO) (7) have been widely used across different data resource projects, which significantly promote standard data annotation. We have developed the Kidney Tissue Atlas Ontology (KTAO) (8) to support kidney tissue knowledge representation. The HuBMAP Common Coordinate Framework Atlas ontology (CCFO) is an application ontology built to support the development of the Human Reference Atlas (HRA) by the HuBMAP project (9). KTAO and CCFO use the same set of reference ontologies, such as CL and Uberon, which support the shared knowledge query and analysis. However, still missing are ontologies to integrate metadata that can support data standardization and integration across domains.

The goal is to create a dynamic integration across different domains (clinical, pathological, and molecular) and multiple data resources that allow us to use structured language to describe an individual’s renal disease. The resulting ontology facilitates the harmonization of language and communication across personas (including pathologists, nephrologists, molecular biologists, bioinformaticians, patients, etc.) and multiple collaborative efforts. KPMP Community Engagement Committee is a workstream focused on understanding patient participants as integral partners in precision medicine research, including understanding experience with barriers to care and needs for inclusive and ongoing engagement in biomedical research. The clinical and associated demographic characteristics (e.g., sex, race, ethnicity, accessibility, language) will be used to create an inclusive format for all individuals. The development and use of the KTAO ontology reflects the inclusive goals and data collection practices of the KPMP project.

This study describes our ontology-based strategy to address the data integration challenge to support kidney precision medicine research. We have developed an integrated precision medicine metadata ontology (PMMO) to collect, map, standardize, and integrate various metadata variables from data resources. PMMO, KTAO, and CCFO are used together to support our kidney precision medicine use case studies. This work demonstrates how ontology-based approaches support data and knowledge integration in precision medicine.

## Methods

Figure 1 illustrates the general project design for data integration across domains. First, we developed PMMO to map, standardize, and integrate metadata variables from multiple resources, including KPMP, HuBMAP, CellxGene, and HCA. HuBMAP and CellxGene only contain single-cell transcriptomic data; notably, the metadata types related to the cell types and the tissues/species/phenotypes of these cell types may differ between HuBMAP and CellxGene. PMMO, however, will represent all cell types and tissues/species/phenotypes. Second, we focus on ontology-based knowledgebase, which represents knowledge with ontologies such as CL, Uberon, HPO, GO, and OPMI. As a related use case, we have identified healthy and kidney disease biomarkers from CCFO and KTAO, respectively. The expression profiles of these biomarker genes from various datasets stored in CellxGene will be queried and analyzed. Since CellxGene does not contain all the metadata of KPMP disease studies (e.g., diabetes phenotype), the metadata ontology will be used to cross-link metadata details in an ontology-standardized manner. The metadata ontology will include sufficient granularity to ensure the cross-integration among different data resources.

**Figure 1.**
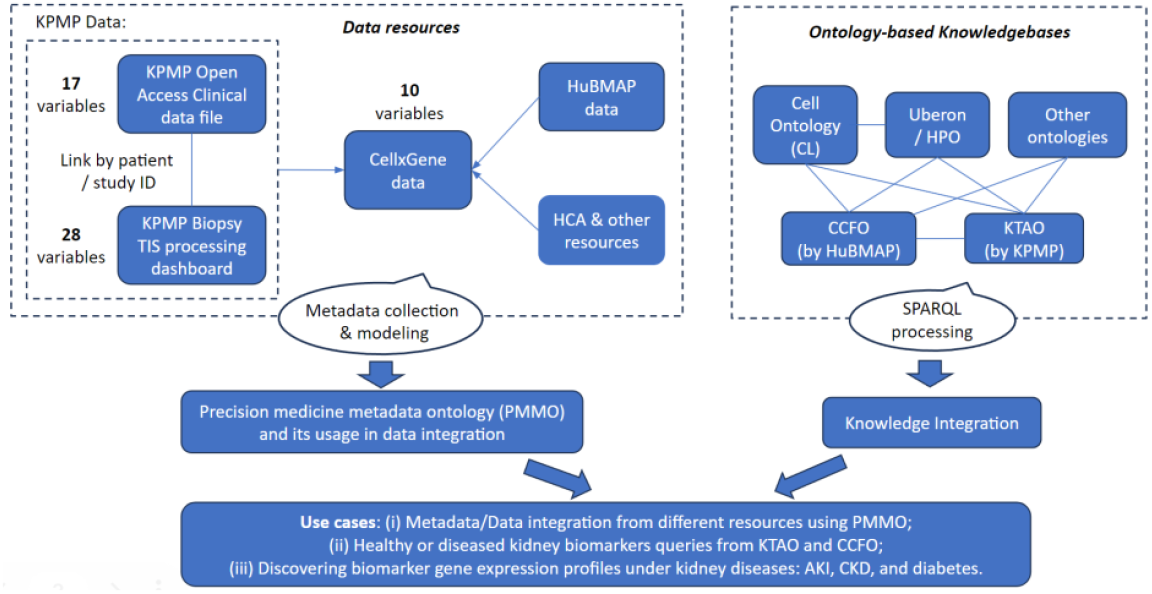
General project design.

### Kidney phenotype annotation and submission to HPO

Starting in 2017, KPMP has organized multiple workshops where nephrologists, pathologists, and ontologists worked together to discuss and apply a high-level framework for annotating, collecting, and proposing new terms or modifications to HPO (7) and underlying ontologies such as Uberon (6). After the internal collection of terminologies and discussion within KPMP, Excel spreadsheets of new terms, modifications, definitions, and proposed hierarchies were prepared and sent to HPO developers. Individual HPO GitHub issue trackers were then generated for individually proposed terms and tagged with “KPMP”. After further adjudication in GitHub and with KPMP investigators, the newly proposed kidney-related HPO terms were added to the HPO. The released terms were then imported back to the KTAO using the tool Ontofox (10).

### Precision Medicine Metadata Ontology (PMMO) – Ontological representation and integration of metadata types from data resources in KPMP and CellxGene

KPMP includes various metadata types associated with clinical, pathological, and molecular aspects (https://www.kpmp.org/available-data). Clinical data were obtained from electronic health records and stored using RedCap (11). Kidney tissue-related data were obtained and stored at the KPMP Tissue Interrogation Sites (TIS) data and pathology data resource. The CellxGene has ten variables (i.e., age, disease, species, tissue, sex, ethnicity, assay, cell type, suspension, donor, and disease) (4). The PMMO ontologically models and represents the KPMP-derived data, the ten common variables (e.g., age, disease) derived from CellxGene, and their semantic relations. Additional new terms and these variable terms were added to enable us to build the semantic relations. Pre-existing ontology terms are reused. The ontology of precision medicine and investigation (OPMI) (8) was the default ontology for the metadata ontology development.

### Kidney disease modeling and analysis with heterogeneous data from multiple resources

Different types of kidney diseases (including AKI, CKD, and others) and their associated pathological features were first modeled using an ontological approach. Then, the data from different resources using KPMP and CellxGene were used to identify, integrate, and analyze the data for the patients related to the modeled kidney diseases.

### Kidney biomarkers collection and representation in KTAO

Gene/protein biomarkers related to kidney cell types and diseases were manually collected from three resources: (i) the KPMP consortium, (ii) our manual literature mining, and (iii) HuBMAP (for reference kidney cell biomarkers). The annotated biomarkers were then added to the KTAO ontology using a standard format. SPARQL (a recursive acronym for SPARQL Protocol and RDF Query Language) queries were generated to extract the biomarker records from HuBMAP CCFO and KPMP KTAO ontologies. The Ontobee SPARQL endpoint (12) was used to generate the query.

### Extraction of kidney biomarker gene expression profiles using CellxGene and KPMP Atlas Explorer

The CellxGene API was used to extract gene expression profiles from specific datasets, and the KPMP Atlas Explorer was utilized for data browsing and analysis. Expression levels for the SPP1 gene were mapped onto relevant cell types in CL using the Graphviz dot language, and the ontology hierarchy was visualized with Graphviz.

This collaborative work involved experts and trainees from different domains. It was impossible for one type of expert to contribute all the domain knowledge. The nephrologists, kidney biologists, pathologists, bioinformaticians, programmers, and ontologists, from experienced professionals to students, met over a year through bi-weekly Zoom calls to discuss and finalize our scientific questions and hypotheses, as well as address data analysis and visualization workflows, ontology modeling, user interfaces, and use case demonstrations. This work also involved collaboration between the KPMP and HuBMAP teams to achieve efficient, interoperable kidney-specific data integration. One graduate student and two undergraduate students also actively participated in this teamwork project.

## Results

### Ontological modeling of kidney diseases and associated pathological phenotypes

We collected hundreds of kidney-specific phenotypes and found many that were not yet in the HPO. Since 2017, our KPMP kidney community has worked closely with HPO developers on the ontological modeling of kidney phenotypes. As of March 16, 2024, HPO had received 329 kidney-specific phenotype terms contributed by KPMP and added most of these terms to HPO. These and other kidney-related HPO phenotype terms were later imported back to the Kidney Tissue Atlas Ontology (KTAO), which aimed to collect and interlink kidney phenotypes and other types of entities (e.g., cells, anatomical entities, biomarkers). By doing so, the KPMP team contributed to the community effort of kidney phenotype collection and modeling for newly developed phenotype ontology models to support KPMP research goals.

A more recent effort is to further model and analyze novel and more specific kidney phenotypes that cross and overlap existing phenotypes. Our KPMP team has been focusing on AKI and CKD. Interestingly, AKI and CKD are not always distinct entities, and overlap or transition of clinical and pathologic characteristics between AKI and CKD can be observed. This definition results in a spectrum of phenotypes (Figure 2). Both AKI and CKD patients may have diabetes and/or hypertension (Figure 2A). For example, patients without any other pathologic conditions may experience AKI alone (i.e., following the use of nephrotoxic antibiotics), but the same AKI phenotype may be superimposed on an underlying diabetic nephropathy phenotype under similar circumstances (diabetic patient with recent use of nephrotoxic antibiotics developing acute renal failure). Similarly, patients with CKD may have characteristics typical of diabetic nephropathy, which is generally a chronic damage phenotype; however, some patients present with a mixed phenotype of chronic active damage, where, in the absence of an obvious trigger, a smoldering acute process occurs hand to hand with the development of chronicity (Figure 2B).

**Figure 2.**
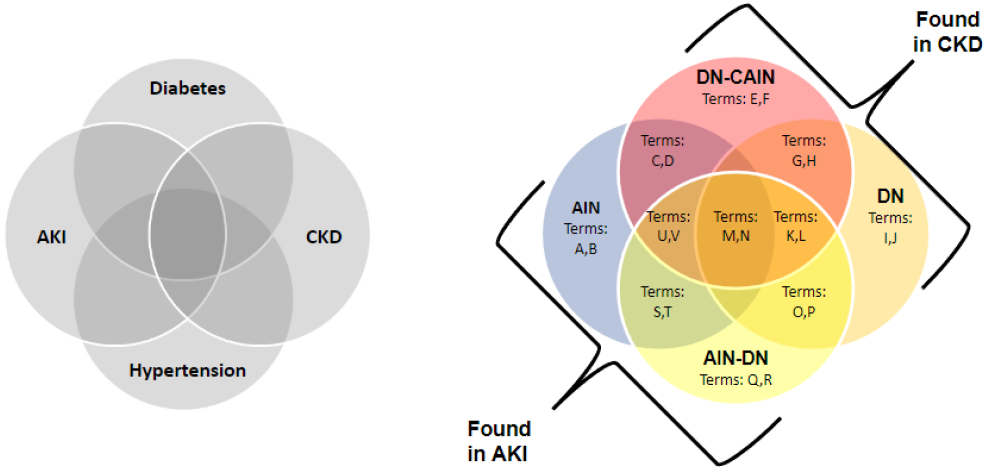
Pathological kidney disease categorization. (A) Venn diagram illustrating how manifestations of diabetes and hypertension may overlap between CKD and AKI. For example, clinical diabetes and hypertension will have terms that are specific for diabetic nephropathy and hypertensive nephropathy (CKD), and terms that overlap with AKI. (B) Venn diagram illustrating how the inclusion of pathology data stratifies patients with diabetic nephropathy into categories manifesting as pure CKD (DN), intermediate phenotypes as diabetic nephropathy with features of chronic active interstitial nephritis (DN-CAIN), and patients with AIN into categories manifesting as pure AKI, or as intermediate phenotypes between AKI and CKD, for example, acute interstitial nephritis superimposed on diabetic nephropathy (AIN-DN).

Our research found that shared phenotypes may have specific features newly identified in different domains. As a result, the overlapping phenotypes may deserve new ontological term representation (Figure 2B). Specifically, we were able to identify two specific disease categories associated with AKI (Figure 2B): Acute Interstitial Nephritis (AIN) and Acute Interstitial Nephritis superimposed on Diabetic Nephropathy (AINs-DN), and two specific disease categories associated with CKD: Diabetic Nephropathy with Chronic Active Interstitial Nephritis (DN-CAIN) and Diabetic Nephropathy (DN).

Many of these kidney disease categories are not yet fully detailed in HPO, and our modeling of these diseases (Figure 2B) is also unique in that it provides a strategy for identifying various disease characteristics indicative of these specific disease categories, thus enabling a more precise definition of disease.

Our current KPMP research focuses on identifying these distinct disease states and their characteristics. One step towards this goal is the work of the KPMP adjudication committee, which routinely conducts a pathology-focused adjudication process that defines disease categories for individual human subjects enrolled in the KPMP study. A descriptor-scoring approach is also used to score different categories of morphologic parameters. After the adjudication-based categorization, these kidney patients are closely monitored to identify unique parameters indicative of their disease categories.

### Integrative KTAO development

As shown in Figure 3, KTAO is an application ontology that serves as the knowledge base of the kidney tissue atlas. As of March 16, 2024, KTAO has 11,882 terms and 175,926 axioms. The KTAO terms include 11,210 classes, 399 object properties, 26 data type properties, 207 annotation properties, and 40 instance terms. In ontology, an axiom represents a relation among terms or entities. Here is an example of an axiom representing the anatomical location of the podocyte (CL_0000653): *‘part of’ some ‘visceral layer of glomerular capsule’* (UBERON_0005751). With so many axioms represented, KTAO serves as its own kidney tissue atlas knowledge base.

**Figure 3.**
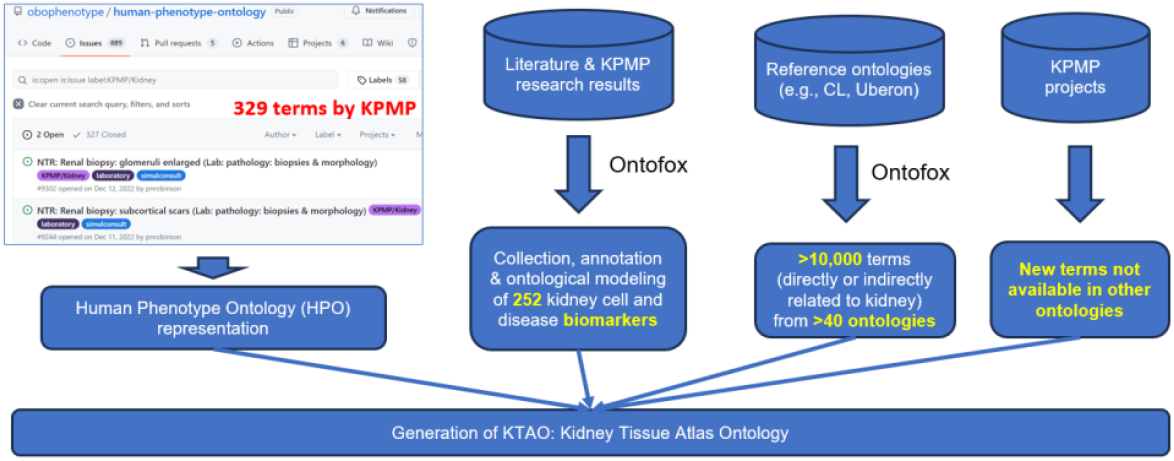
KTAO development workflow and statistics. As of February 26, 2024, our KPMP team had contributed to HP 329 terms, and 327 of them had been added to HPO and then imported back to KTAO. We also collected 252 kidney-related biomarkers and represented them in KTAO. KTAO has also imported more than 10,000 terms from over 40 existing ontologies. Furthermore, KTAO generates KTAO-specific terms that are not available in other ontologies.

KTAO reuses existing ontologies such as CL, HPO, and Uberon. KTAO imports terms from 32 ontologies with at least ten terms included (https://ontobee.org/ontostat/KTAO). Table 1 lists the statistics of terms in KTAO.

**Table 1.** KTAO statistics of terms, including those imported from other ontologies and KTAO-specific terms.

| Ontology used in KTAO (selected) | No. of terms in KTAO |
| --- | --- |
| Cell Ontology (CL) | 386 |
| Human Phenotype Ontology (HPO) | 960 |
| Uber-anatomy ontology (Uberon) | 1,171 |
| Gene Ontology (GO) | 1,225 |
| Protein Ontology (PR) | 316 |
| Ontology of Genes and Genomes (OGG) | 483 |
| Ontology of Precision Medicine and Investigation (OPMI) | 864 |
| KTAO-specific (with KTAO namespace) | 53 |
| Total in KTAO (version 1.0.121, 2024-03-16) | 11,882 |

### PMMO development and application for precision medicine metadata integration

To efficiently analyze heterogeneous data from different data resources, it is critical to integrate the data types at the metadata level. However, such a task is often challenging. As illustrated in Figure 4, we focused on the task of integrating three specific metadata types: (i) 18 KPMP clinical metadata types focused on clinical data, (ii) 29 KPMP biopsy TIS metadata, and (iii) 10 CellxGene metadata types. Our PMMO systematically models and represents these metadata types in an integrative semantic framework.

**Figure 4.**
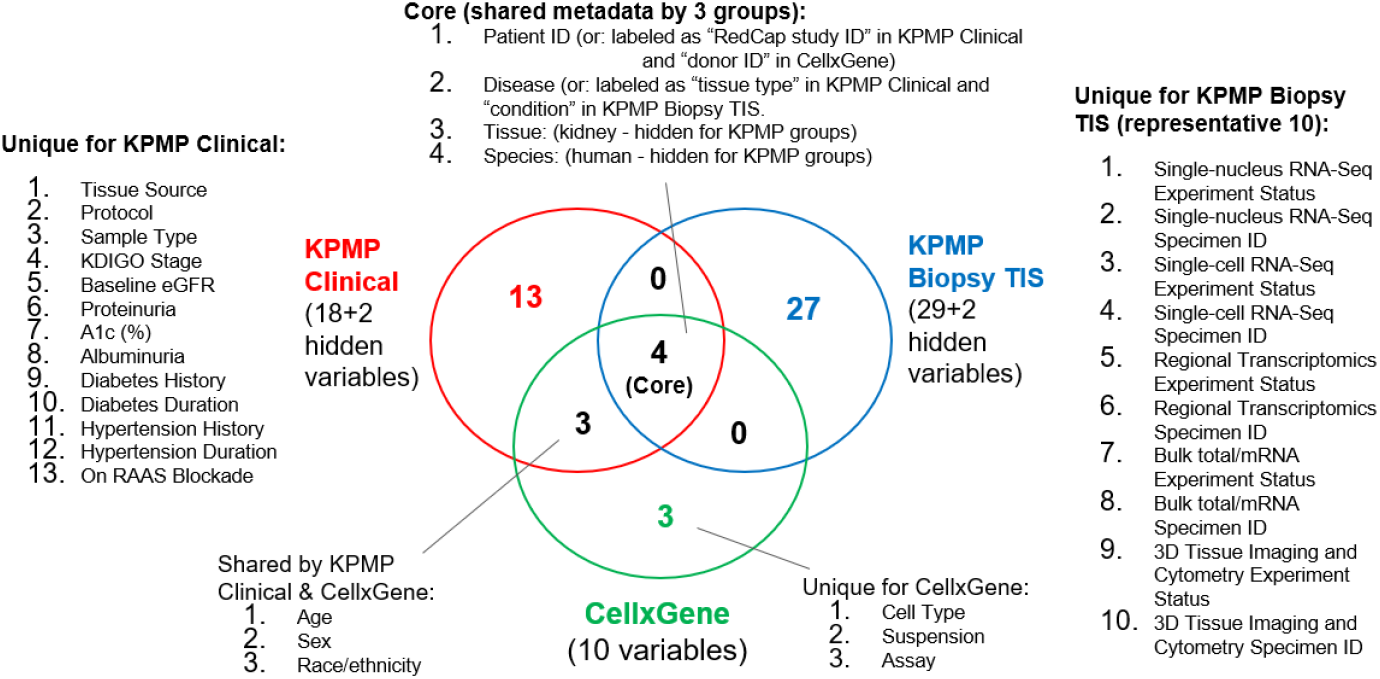
Venn Diagram of metadata variables from three specific data resources.

We realized that even with small numbers of metadata elements, significant issues arise in our integration and modeling of the three metadata sets (Figure 4). For example, the core data type ‘patient ID’ is defined using different labels: ‘patient ID’ in the KPMP Biopsy TIS data, ‘RedCap study ID’ in KPMP Clinical data, and ‘donor ID’ in CellxGene data. While these terms are used in different scenarios, we found that they refer to the same IDs and can be used as the most important metadata elements to interlink different data resources. Another example is the representation of ‘disease’ in different data resources: ‘disease’ in CellxGene data, ‘tissue type’ in KPMP clinical data, and ‘condition’ in KPMP Biopsy TIS data (Figure 4). In addition to ‘tissue type’, the KPMP clinical data file has another tissue-related metadata element called “Tissue Source”. Here, “tissue type” refers to specific disease information (e.g., AKI, CKD, and healthy), and “tissue source” refers to where to obtain the tissue (e.g., KPMP TIS). Mapped to the “Tissue” and “Species” in CellxGene, the two KPMP datasets have two hidden values: kidney for “Tissue” and human for “Species”. Therefore, the three metadata sets share four data types: patient ID, disease, tissue, and species, which have been properly identified and modeled in PMMO.

In addition to the core terms shared by all three metadata sets, we found many metadata elements unique to each set or shared by only two of them (Figure 4).

The PMMO representation reuses terms from reference ontologies. We first mapped the different metadata elements to terms from reference ontologies and then integrated them into one ontology format. The OPMI is used as the default ontology for the PMMO ontology development.

An important use case of the PMMO is in integrating information from different sources. Guided by the PMMO modeling, we could trace the kidney precision medicine data across different data resources using KPMP/CellxGene web browsers (Figure 5) or API. For example, using the KPMP repository report program, we are able to identify the records of an anonymized human subject with the patient ID of “27-10039” (Figure 5A). This human subject is a CKD patient (female, white, 60-69 years old) with a diabetes history of 30-40 years and a hypertension history of 30-40 years (Figure 5B). The expression profiles of the gene *SLC5A2* (solute carrier family 5 member 2, commonly known as SGLT2) are shown in KPMP spatial transcriptomics format (Figure 5C) or KPMP gene expression format (Figure 5D). Furthermore, CellxGene also provides specific profiles for this human donor with the donor ID as the same as the KPMP patient ID (Figure 5E). The proposed PMMO will include the linkage information to interlink datasets between different datasets.

**Figure 5.**
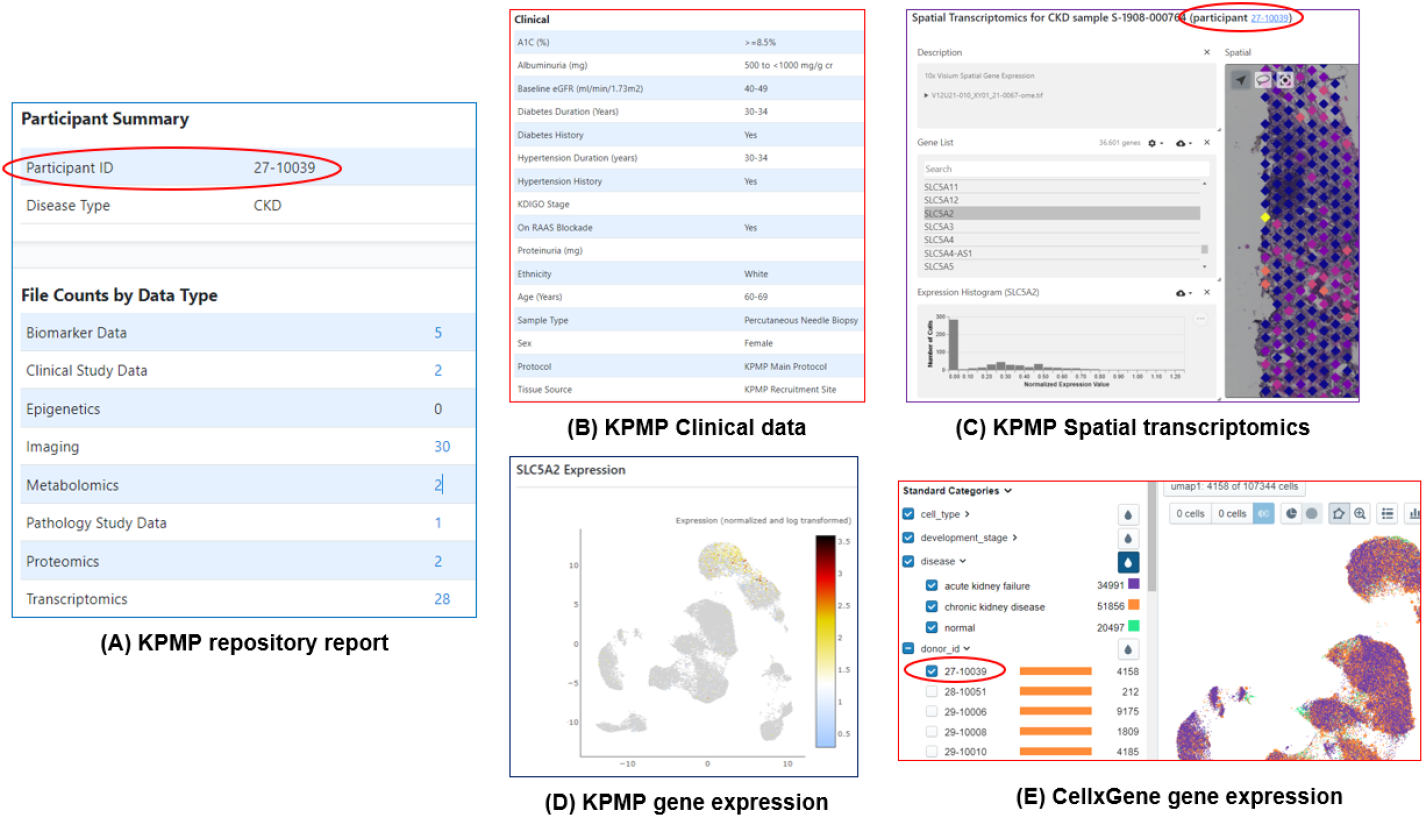
Integrative exploration of the records of an anonymized KPMP human subject (patient ID: 27-10039) using KPMP or CellxGene web programs. (A) Basic KPMP information of records associated with this patient. (B) Clinical data. (C) KPMP spatial transcriptomics related to the example gene SLC5A2. (D) KPMP gene expression profile for SLC5A2. (E) CellxGene record whose donor ID equals to KPMP patient ID.

Using the same approach as laid out above, we can search for all the datasets across the different sources, including KPMP clinical data, KPMP biopsy TIS data, and CellxGene. KPMP systematically collects and analyzes the records and kidney biopsy samples from well-studied human subjects with AKI, CKD, or reference healthy controls. The related cell gene expression data from the KPMP studies have also been submitted to CellxGene. It is important to note that different data sources use the same IDs, although these IDs have different labels (patient ID, RedCap study ID, or donor ID) (Figure 4). Different databases also have different variables. For example, the CellxGene source has ten variables (e.g., sex, age, disease), some of which are unique or shared with other databases (Figure 4). These variables can be analyzed using the web browsers or the CellxGene API.

### KPMP kidney gene/biomarker collection, annotation, and analysis

Our research identified 529 kidney gene biomarkers (Figure 6A). These biomarkers include 192, 26, and 19 gene biomarkers unique to healthy kidney or kidney cell types, CKD, or AKI, respectively. In addition, these biomarkers include three sets shared by two groups: 3 shared by healthy and CKD groups, 5 shared by healthy and AKI groups, and 14 shared by AKI and CKD groups (Figure 6A). These biomarkers were ontologically represented in KTAO.

**Figure 6.**
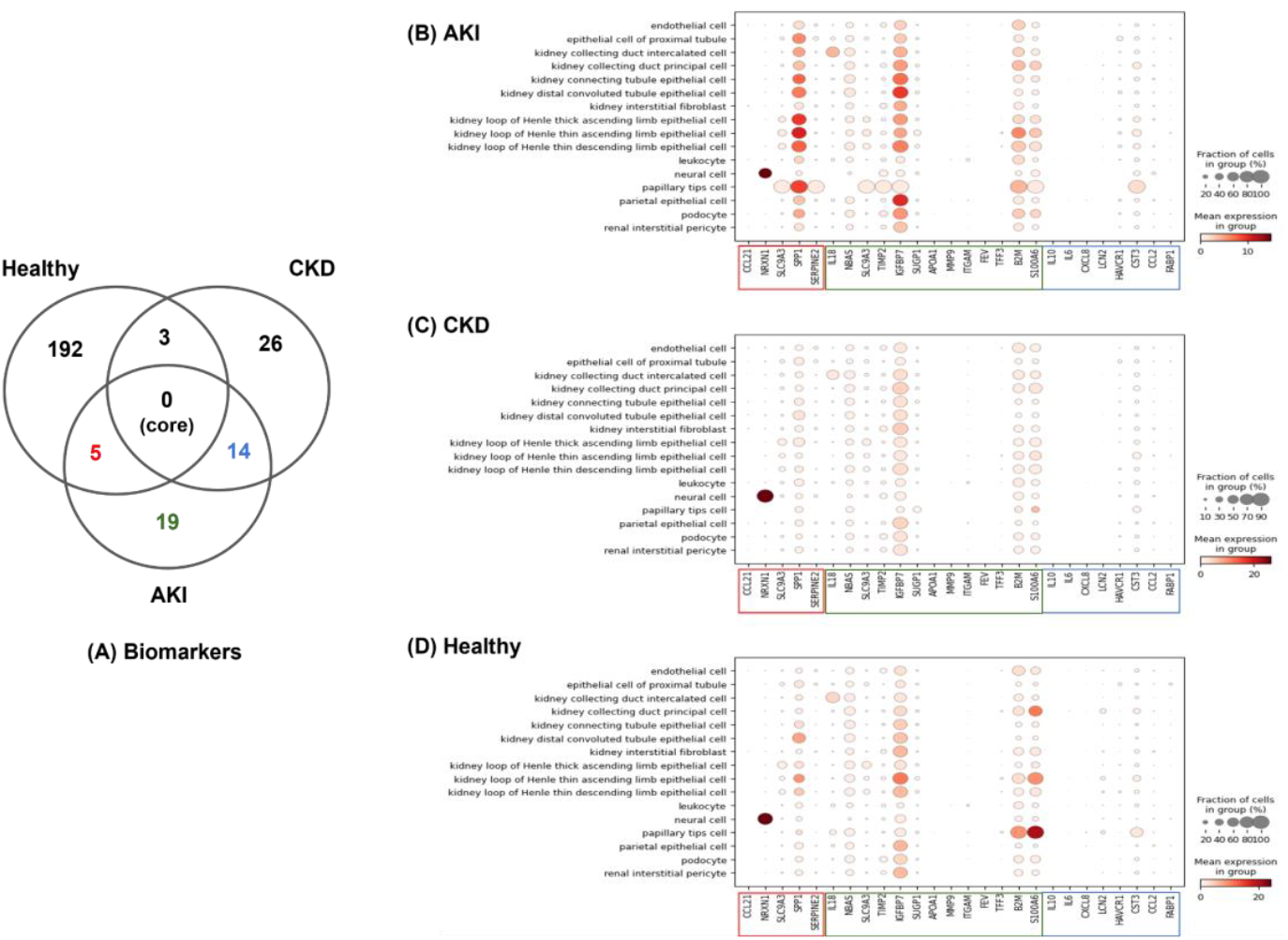
Kidney gene biomarker analysis. (A) Kidney biomarkers collected in KTAO. (B-D) The gene expression profiles of 26 AKI biomarker genes using the KPMP study labeled “Single-nucleus RNA-seq of the Adult Human Kidney (Version 1.0)” (Dataset_id in CellxGene: 07854d9c-5375-4a9b-ac34-fa919d3c3686). (B) AKI patient condition. (C) CKD patient condition. (D) healthy condition. CellxGene API was used to generate the (B-D) results. The colors in (A) match the same colored boxes in (B-D).

Kidney biomarkers were also analyzed using different resources. Specifically, we focused on the 38 AKI biomarkers, as shown in Figure 6A, using the KPMP datasets described in a *Nature* paper (14). The dataset is also stored in the CellxGene with a specific Dataset_id. Note that only 26 out of these 38 AKI biomarker genes had data in the CellxGene dataset. These 26 genes include five genes shared by healthy and AKI groups (red color), eight genes shared by CKD and AKI groups (blue color), and 13 gene markers unique to AKI (green color) (Figure 6B-D). Our results show that some biomarkers can be confirmed to be good markers, but some not. These biomarkers have different profiles.

While Figure 6 provides a detailed presentation of 26 gene biomarkers’ gene expression profiles among 16 kidney cell types, these 16 cell types have additional relationships that are not observable in the figure. To address relationships between cell types, we mapped these cell types to the Cell Ontology (CL) and used the CL hierarchy to identify the relations among these cell types (Figure 7). By doing so, we were able to obtain more insights. For example, with the help of the CL hierarchical structure, we were able to find that SPP1 was highly expressed in the high-level class term “nephron tubule epithelial cell” (CL_1000494) in the healthy controls and AKI patients (but not in CKD patients), esp. in its subclass of “kidney loop of Henle ascending limb epithelial cell” (CL_1000849), which includes two specific cell types: “kidney loop of Henle thin ascending limb epithelial cell” (CL_1001107) and “kidney loop of Henle thick ascending limb epithelial cell” (CL_1001106). Furthermore, the SPP1 expression profile in the “kidney loop of Henle thick ascending limb epithelial cell” appears to be distinct among three disease statuses (healthy, AKI, and CKD) (Figure 7). Since SSP1 protein functions as a cytokine, these differential expression patterns can help us identify changes in signaling functions across cell types and the different regions of the nephron. If these changes in signaling lead to changes in metabolism and/or transport, it is reasonable to hypothesize these changes could be reflected in new metabolites in the urine that could serve as biomarkers.

**Figure 7.**
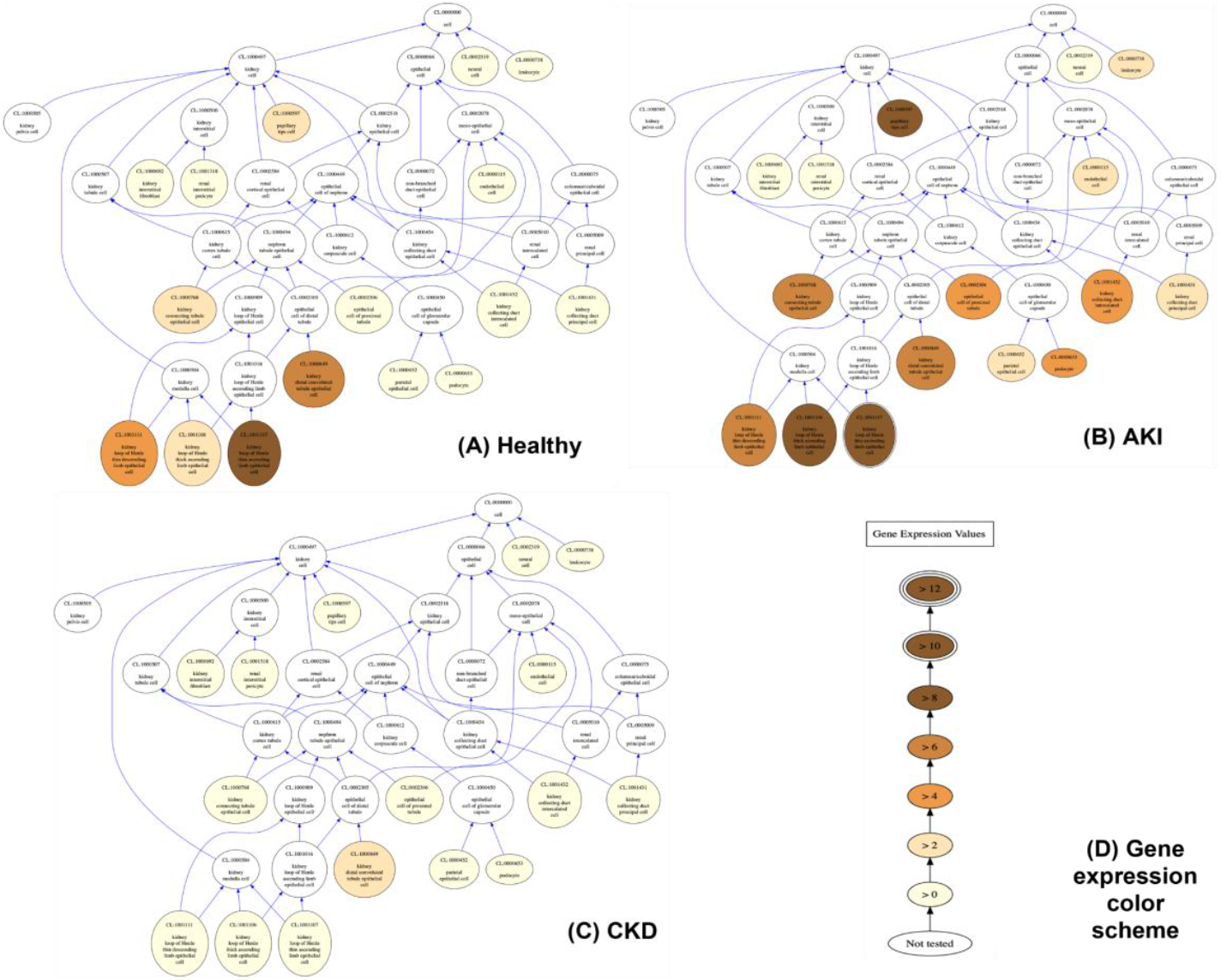
Ontology-based analysis of SPP1 gene expression profiles in 16 different cell types. Cell types arranged in the Cell Ontology (CL) hierarchy for kidney cell types were colorized based on their measured expression of the SPP1 gene, with darker shades indicating higher expression levels based on the log_10_ of the expression value.

## Discussion and Conclusions

This study describes several substantive advances in providing a semantic framework for the kidney atlas. First, we are developing an approach for modeling kidney-specific histological and pathological phenotypes, leading to a collaborative effort of adding more than 300 kidney-specific phenotype terms to the HPO ontology. We also report our continued identification and ontological modeling of many precise kidney disease subcategories (Figure 2). Second, KTAO was applied to systematically integrate various types of kidney-related knowledge at the ontological level. For example, KTAO imports terms and axioms related to kidney cells, anatomic entities, and phenotypes from CL, Uberon, and HPO, respectively. It also collects and annotates various kidney-related biomarkers from KPMP and HuBMAP collections and our literature mining. Third, we have developed the PMMO and applied it to seamlessly integrate data from different data resources (e.g., KPMP clinical data resource, KPMP tissue interrogation and Omics data resource, and CellxGene resource). Fourth, we have applied our ontological modeling and analysis strategy for kidney disease use case studies, focusing on the AKI and CKD disease characteristics and gene biomarker profile analysis using different resources. We provided a descriptive architecture to support downstream applications for quantifiable data labels.

We find challenges in systematically modeling and analyzing many specific kidney disease sub-categories. For example, our KPMP disease-focused analyses enumerated many kidney disease subcategories by overlapping higher-level disease categories such as AKI, CKD, and diabetes (Figure 2). These disease subcategories appear to have unique features, and our precision medicine approaches provide a feasible strategy to find these unique features by profiling and integrating clinical, pathological, cellular, and molecular characteristics related to an increasing pool of human subjects from various backgrounds and identities with these specific diseases. A collaborative effort involving kidney clinicians, biologists, pathologists, ontologists, and bioinformaticians is required to further categorize and identify unique features of these disease subcategories in multiple domains.

To address the challenge of integrating multi-domain features, we propose to develop KTAO further or separately develop a new integrative application ontology tentatively called “Pathomics Ontology” (or Histology Descriptors and Pathomics Ontology - HDPO). The KPMP analysis of tissue includes a variety of methodologies, including i) disease adjudication, which takes into account the clinical data and the overall histology, ii) descriptor scoring, which is a detailed visual assessment and scoring of all kidney tissue compartments, and iii) the pathomic features, which uses computational image analysis to quantify morphometric and textural tissue characteristics extracted from automatically segmented functional tissue units. The HDPO will combine the pathology phenotyping and descriptor scoring with the pathomic features and omics data. The goal is to identify unique features associated with specific pathological phenotypes at the cell, structure, and biopsy level. This proposed ontology will reuse terms from existing ontologies (e.g., CL, Uberon, and HPO) and incorporate many other concepts, such as histological descriptors, pathomic features, and omics-related terms from spatial transcriptomics, with the goal of better supporting systematic and integrative modeling and analysis of newly identified disease or phenotype subcategories and unique features associated with these subcategories. The KPMP descriptor scoring and computational image analysis processes provide a solid base and model for such ontology development and application.

Data from different resources can be integrated in different ways. At the level of knowledge (a special type of annotated data), an efficient way is to incorporate the learned knowledge into ontologies and knowledge graphs and then use programming languages such as SPARQL to perform queries and analyses. For example, we successfully developed SPARQL scripts to query and analyze biomarker knowledge from the KTAO and HuBMAP CCFO. At the instance data level, we posit that metadata integration is critical for data integration from different sources. Our research has identified the critical need for metadata ontology to integrate different precision medicine-related data sources, which led to our initiation of the development of PMMO metadata ontology. We demonstrate applications of PMMO in our kidney precision medicine research. This type of metadata ontology and its associated methods can also be applied to study other organs and diseases, including the heart, liver, and vasculature.

Our ontology-based integrative approaches, using knowledge from multiple domains, can drive the discovery of new biomarkers. While hundreds of kidney gene biomarkers have been found, our research focused on data integration has found that not all these biomarkers are equally expressed in kidney cells. Using our ontology-based methodology, it should be possible to identify the key cell types involved in the different kidney disease states and find a small number of gene biomarkers that work well at individual cell levels.

In the future, we expect to maintain and extend our mappings to more scenarios and use cases. We will also develop our community-based Precision Medicine Metadata Ontology (PMMO) further and apply it to support integrative studies across different domains and projects, such as KPMP, HuBMAP, HCA, CellxGene, etc. We expect that such an ontology-based strategy will significantly improve computer-understandable data standardization and integration, leading to advanced data analysis and scientific question exploration.

## Acknowledgments

The KPMP is funded by the following grants from the NIDDK: U01DK133081, U01DK133091, U01DK133092, U01DK133093, U01DK133095, U01DK133097, U01DK114866, U01DK114908, U01DK133090, U01DK133113, U01DK133766, U01DK133768, U01DK114907, U01DK114920, U01DK114923, U01DK114933, U24DK114886, UH3DK114926, UH3DK114861, UH3DK114915, UH3DK114937. Human Reference Atlas (HRA) research and development is funded by the NIH Common Fund through the Office of Strategic Coordination/Office of the NIH Director under awards OT2OD033756 and OT2OD026671, by the Cellular Senescence Network (SenNet) Consortium through the Consortium Organization and Data Coordinating Center (CODCC) under award number U24CA268108, and by the NIDDK under awards U24DK135157 and U01DK133090. The work has also been supported by the Kidney Precision Medicine Project grant U2CDK114886, and the NIH National Institute of Allergy and Infectious Diseases (NIAID), Department of Health and Human Services under BCBB Support Services Contract HHSN316201300006W/HHSN27200002, and HuBMAP U54 DK134301.

## Notes

https://github.com/KPMP/KTAO

https://www.kpmp.org/available-data

https://cellxgene.cziscience.com/collections/bcb61471-2a44-4d00-a0af-ff085512674c

https://github.com/OPMI/opmi

### Competing Interest Statement

The authors have declared no competing interest.

### Summary of Updates

The summary of the revision is below: - Figure 2 has been updated. - Author list updated with one author removed as the same author suggested to be removed. - Editorial changes in the manuscript.

